# Amoxicillin Worsens Mortality from Viral Respiratory Tract Infection

**DOI:** 10.64898/2026.10.27.762283

**Authors:** Marika Orlov, Steven C. Shaw, Sara N. Stoner, Sam Fulte, William J. Janssen, Sarah E. Clark

## Abstract

Influenza infections are common and result in substantial mortality, with secondary bacterial pneumonias contributing to worse outcomes. To prevent mortality associated with bacterial co-infections, antibiotics are often prescribed in the absence of microbiological data. Currently, there are conflicting society recommendations on antibiotic utilization in viral respiratory tract infections as we lack prospective studies to guide treatment. We developed an animal model to study the effects of antibiotic exposure during influenza infection. Our results suggest that amoxicillin started during influenza A infection increases mortality from influenza A and increases severity of secondary bacterial pneumonia, and this is more pronounced in males.

## Introduction

Influenza respiratory tract infections are responsible for millions of outpatient visits, hundreds of thousands of hospitalizations and tens of thousands of deaths annually. Secondary bacterial pneumonia is an important contributor to severe influenza-associated disease and mortality (1), with *Streptococcus pneumoniae* among the most common bacterial pathogens identified during influenza-associated co-infection (2, 3). For this reason, the most recent American Thoracic Society guidelines recommend empiric antibacterial coverage for higher risk individuals for community acquired pneumonia while gathering microbiological data (4). The Infectious Diseases Society of America does not endorse these guidelines and cautions against antibacterial coverage in patients that test positive for a viral respiratory tract infection until there is clear evidence of bacterial co-infection (5, 6). Currently, there is a lack of prospective studies to guide appropriate treatment.

This uncertainty is particularly important because antibiotic exposure itself may influence the course of viral infection and susceptibility to subsequent bacterial pneumonia. A large cohort study showed that patients prescribed antibiotics for influenza infection in the outpatient setting had an associated increase in hospitalizations and longer hospital stays (7). Similarly, in hospitalized patients with confirmed respiratory viral infections, antibiotic treatment initiated later during hospitalization was associated with increased mortality (8). Experimental studies provide a potential biological basis for these observations: antibiotic-mediated disruption of the gut microbiota impairs pulmonary antibacterial immunity (9), while influenza independently suppresses host defense against secondary bacterial pneumonia (10). Together, these findings raise the possibility that antibiotic exposure during influenza may further impair pulmonary infection defense. However, whether treatment with antibiotics initiated *after* the onset of influenza alters viral disease outcomes or subsequent bacterial pneumonia outcomes remains unclear. We therefore sought to establish a clinically relevant animal model to study the effects of antibiotic exposure in the setting of viral respiratory tract infection.

Although antibiotic exposure has been incorporated into murine models of influenza-associated secondary bacterial pneumonia, most studies initiate antibiotics before influenza infection. We developed a model to test the clinical scenario of antibiotic treatment during influenza infection to determine the impact of antibiotics on viral infection and secondary bacterial pneumonia. Unexpectedly, we observed significantly increased mortality in mice that were started on amoxicillin during influenza infection, even without a secondary bacterial infection.

## Methods

### Mice

All studies were conducted with the approval of the Institutional Animal Care and Use Committees at the University of Colorado. Wild type C57BL/6 mice were used. Equal numbers of males and females were used

### Influenza Infections

Influenza A virus (IAV, mouse adapted strain x31) stocks were provided by Dr. Jenna Guthmiller, University of Colorado Anschutz. 7-8 weeks old mice were anesthetized with inhaled isoflurane and then intranasally infected with IAV at either a “high” dose of 5x10^3^ plaque forming units (PFUs) or a “low” dose of 1x10^3^ PFUs. Survival was monitored in mice that received high dose IAV infection.

### Amoxicillin treatment

Three days after inoculation with IAV, half the mice received amoxicillin (0.26 mg/ml) *ad libitum* in their drinking water for 5 days while the other half received control, unmedicated water.

### *Streptococcus pneumoniae* (Spn) infections

In the low dose IAV infection group, mice were infected with *Streptococcus pneumoniae* (Spn) via oropharyngeal aspiration 11 days after IAV infection. The Spn strain used for these experiments was serotype 2 strain D39 (11), which is an invasive strain currently targeted in the Pneumovax 23 vaccine. Spn inoculum was grown in Todd Hewitt Broth (with 5% Yeast Extract (BD Bacto™) from frozen stocks at 37 °C with 5% CO2 with no shaking to mid-log phase (O.D. range: 0.3-0.8) and centrifuged at 20,000 x g for 10 min at room temperature followed by resuspension in PBS for infections (3x10^6^ colony forming units (CFUs) in 50 µl/mouse). Amoxicillin was removed from the drinking water for 3 days prior to Spn infections. CFU burdens were assessed 24 hours after infection. Briefly, mice were euthanized and bronchoalveolar lavage (BAL) was collected with 1 mL EDTA/PBS. Lungs and spleens were harvested and subsequently homogenized. BAL fluid, homogenized lungs and spleens were serially diluted for Spn CFU enumeration on selective media.

### Statistics

Significance for survival was assessed using Kaplan-Meyer survival curves and a Mann-Whitney test was used to assess significant differences in bacterial burdens.

## Results

### Amoxicillin increases mortality from viral respiratory tract infections

Male and female adult mice were infected with a high viral inoculum of IAV (5x10^3 PFU) via the intranasal route. Three days later, half the mice were started on amoxicillin water, while the other half were continued on control water. The amoxicillin water was continued for a total of 5 days. Amoxicillin was chosen as it is the most commonly prescribed antibiotic and results in gut microbiome dysbiosis without complete microbiome depletion (12). Survival was monitored in these mice. Surprisingly, mortality was significantly increased in amoxicillin treated animals (p=0.04, Figure 1A). This difference was driven by male mice, all of whom died with amoxicillin treatment (p=0.01, Figure 1B). In contrast, there were no significant mortality differences in female mice regardless of amoxicillin treatment (p=0.77, Figure 1C). These data suggest that antibiotics started *during* influenza A infection alter the course of viral infection and contribute to increased mortality in the absence of a secondary bacterial infection and that this effect is more apparent in males.

**Figure 1:**
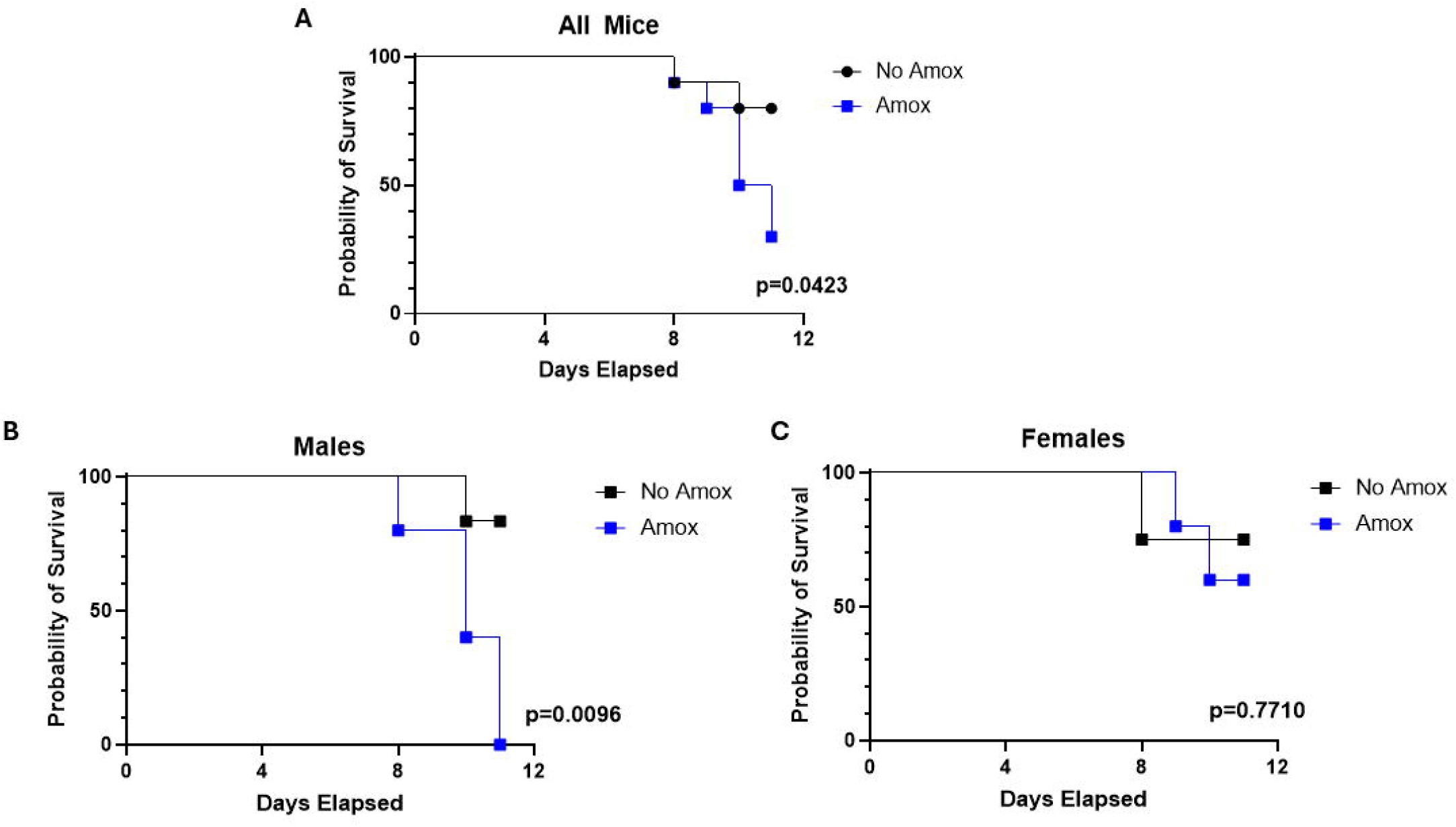
Amoxicillin increases mortality from viral respiratory tract infection. Wild-type mice were intranasally infected with influenza A virus (x31, 5x10^3^). Three days later, mice were treated with amoxicillin (blue squares) or regular water (black circles) *ad libitum* for 5 days. Survival curves shown for all mice (A), only male mice (B), and only female mice (C). N=10 mice/group, 2 pooled experiments. N=10 mice/group, 2 pooled experiments. Kaplan-meyer survival curve, Mantel-Cox Test for significance.

### Amoxicillin increases bacterial burdens in secondary Spn pneumonia

Next, we assessed severity of post-viral bacterial pneumonia with and without amoxicillin treatment. In these experiments, male and female adult mice were infected with a lower inoculum of influenza A virus to allow for secondary bacterial challenge. As before, half the mice received a five-day course of amoxicillin starting 3 days after IAV inoculation. On day 11 post-IAV inoculation, mice were challenged with Spn. Tissues were harvested to measure bacterial burdens 24 hours later.

More mice had detectable burdens of Spn in the spleen in the amoxicillin treated group (10 out of 19 mice, 53%) compared to the mice that received non-medicated water (4 out of 17 mice, 24%, p=0.05, Figure 2A). This suggests an increase in systemic Spn infection in mice treated with amoxicillin despite no difference in overall Spn burdens in the lungs (BAL and Lung CFUs, Figure 2A). When males and females were assessed separately, there was evidence of increased Spn burden in the lungs of male mice (Figure 2B), but not female mice (Figure 2C). These data demonstrate that antibiotics increase Spn dissemination in post viral bacterial pneumonia and that bacterial burdens in the lungs are more pronounced in males with antibiotic treatment.

**Figure 2:**
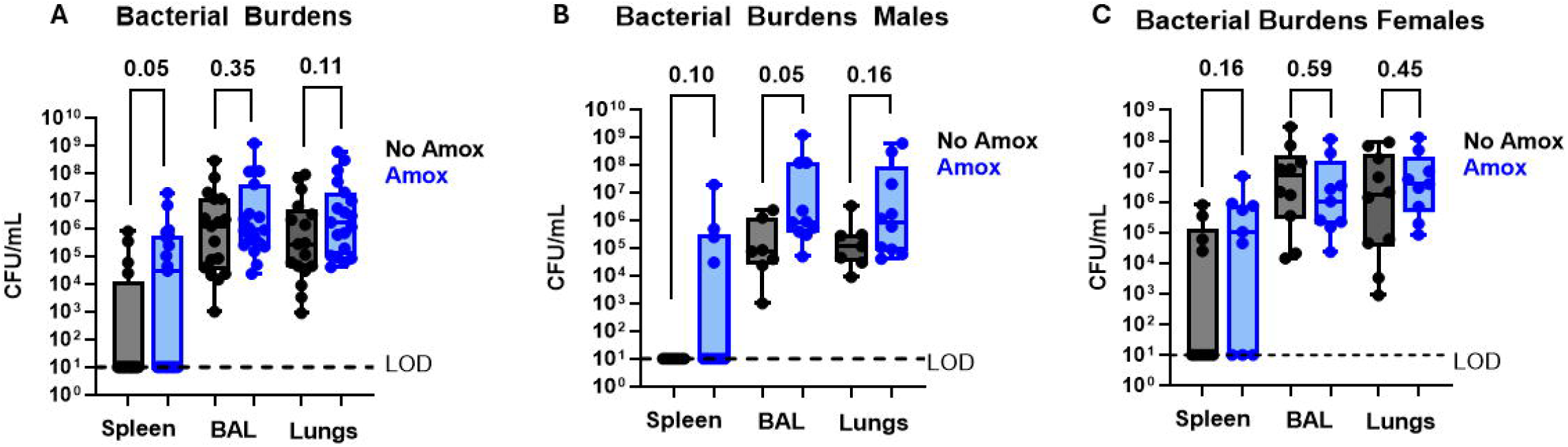
Amoxicillin increases systemic infection with secondary bacterial pneumonia. Wild type mice were intranasally infected with influenza A virus (x31, 1x10^3^). Three days later, mice were treated with amoxicillin (blue circles) or regular water (black circles) *ad libitum* for 5 days. All mice were switched to regular water for three days and then infected with *Streptococcus pneumoniae* (Spn) (3x10^6^ colony forming units (CFUs)) via oropharyngeal aspiration on day 11 post-IAV inoculation. 24 hours later, CFU burdens in the spleen, BAL and lungs were assessed in all mice (A, N=17-19 per group), only male mice (B, N=7-10 per group), and only female mice (C, N=9-10 per group). LOD: Limit of detection. 3 pooled experiments, Mann-Whitney test used to determine significance between groups for each tissue type.

## Discussion

Our data show that administration of amoxicillin after the onset of severe viral pneumonia increases mortality in male mice and impairs host defense against secondary bacterial pneumonia. Previous studies have shown that prolonged microbiota depletion prior to influenza impairs antiviral immune responses and increases pulmonary viral burden (13), while broad-spectrum antibiotic exposure beginning one week prior to influenza infection impairs antibacterial immunity during subsequent bacterial pneumonia (14). Our findings extend these observations by demonstrating that a short course of a single antibiotic initiated *during* influenza infection can alter both viral and post-viral bacterial pneumonia outcomes. This timing more closely reflects the clinical scenario in which antibiotics are prescribed after patients present with a viral respiratory infection. Importantly, these effects occur with an amoxicillin regimen that induces gut dysbiosis (12) without the profound microbial depletion associated with prolonged courses of multiple broad-spectrum antibiotics commonly used in animal studies.

Another significant finding is that male and female mice respond differently to amoxicillin exposure. Although sex differences in antibiotic-associated adverse effects have been reported (15), whether antibiotic exposure can differentially influence mortality or susceptibility to subsequent infection between males and females has not been established. This represents an important knowledge gap, as antibiotics are commonly prescribed across sexes with the assumption that their effects are broadly similar. Our findings raise the possibility that antibiotic exposure may have consequences that extend beyond traditional adverse effects to influence sex-dependent survival and vulnerability to subsequent infection. Further research is therefore needed to determine whether sex should be considered when evaluating the risks and benefits of commonly used antibiotics.

An important aspect of our findings is that amoxicillin altered outcomes both during primary viral infection and following secondary bacterial challenge. Increased mortality during influenza infection in the absence of secondary bacterial infection suggests that antibiotic exposure may alter the host response to an ongoing viral infection, rather than acting solely by increasing susceptibility to subsequent bacterial pneumonia. While influenza itself creates a period of impaired antibacterial host defense and increased susceptibility to secondary bacterial pneumonia, our findings suggest that antibiotic exposure during this period may further compromise the ability of the host to respond to subsequent bacterial infections.

Our study has several limitations. The mechanisms responsible for this effect are yet to be determined. We evaluated a single antibiotic and experimental influenza model, and the extent to which these findings generalize to other antibiotics, respiratory viruses, or bacterial pathogens remain unknown.

Finally, although initiating amoxicillin during influenza infection more closely reflects clinical antibiotic use during viral respiratory illness, studies in human populations will be necessary to determine whether antibiotic exposure during influenza similarly influences disease severity or susceptibility to secondary bacterial pneumonia, though retrospective human studies do support this possibility (7, 8).

Together, our findings identify adverse effects of antibiotic exposure initiated during an established viral respiratory infection and support further investigation in human populations. These results do not diminish the importance of appropriate antibacterial treatment when bacterial co-infection is present but suggest that the potential consequences of unnecessary antibiotic exposure during viral respiratory infections may extend beyond antimicrobial resistance.

